# A Highly Efficient and Faithful MDS Patient-Derived Xenotransplantation Model for Pre-Clinical Studies

**DOI:** 10.1101/265082

**Authors:** Yuanbin Song, Anthony Rongvaux, Ashley Taylor, Tingting Jiang, Toma Tebaldi, Kunthavai Balasubramanian, Arun Bagale, Yunus Kasim Terzi, Rana Gbyli, Xiaman Wang, Jun Zhao, Nikolai Podoltsev, Mina Xu, Natalia Neparidze, Ellice Wong, Richard Torres, Emanuela M. Bruscia, Yuval Kluger, Markus G. Manz, Richard A. Flavell, Stephanie Halene

## Abstract

Comprehensive preclinical studies of Myelodysplastic Syndromes (MDS) have been elusive due to limited ability of MDS stem cells to engraft current immunodeficient murine hosts. We developed a novel MDS patient-derived xenotransplantation model in cytokine-humanized immunodeficient "MISTRG" mice that for the first time provides efficient and faithful disease representation across all MDS subtypes. MISTRG MDS patient-derived xenografts (PDX) reproduce patients' dysplastic morphology with multi-lineage representation, including erythro- and megakaryopoiesis. MISTRG MDS-PDX replicate the original sample's genetic complexity and can be propagated via serial transplantation. MISTRG MDS-PDX demonstrate the cytotoxic and differentiation potential of targeted therapeutics providing superior readouts of drug mechanism of action and therapeutic efficacy. Physiologic humanization of the hematopoietic stem cell niche proves critical to MDS stem cell propagation and function *in vivo*. The MISTRG MDS-PDX model opens novel avenues of research and long-awaited opportunities in MDS research.

## INTRODUCTION

Myelodysplastic Syndrome (MDS) is a group of heterogeneous disorders of the hematopoietic stem cell characterized by recurrent genetic aberrations in genes of essential pathways, including transcription factors, epigenetic regulators, cohesin complex genes, DNA repair genes, and key factors of the spliceosome (^1,2^ and reviewed in ^3^).

Long-term hematopoietic stem cells (HSCs) cannot be expanded in culture and MDS cell lines do not exist, creating an unmet need for *in vivo* models of MDS. Xenotransplantation of primary human MDS stem cells into currently available immunodeficient mice, such as *NOD-scid Il2rg^-/-^* (NSG), has demonstrated limited success with poor and transient engraftment, skewing towards the lymphoid lineage, and engraftment mostly restricted to the injected tibial bone when aided by co-injection of human mesenchymal stem cells (MSC) ^4-7^. Human cytokines provided by constitutive, transgene-driven expression in the NSG-SGM3 model (overexpressing human stem cell factor (SCF), granulocyte-monocyte-colony stimulating factor (GM-CSF), and interleukin-3 (IL3) from a CMV-promoter), improve myeloid differentiation and cellular proliferation, yet stem cell maintenance is impaired ^8-11^. This limitation is overcome transiently by co-injection of autologous human MSC ^12^ or by creation of an ossicle from human MSC that provides an improved human stem cell environment ^13^. These latter two approaches have limited applicability in pre-clinical studies that require a highly efficient, high-throughput approach.

We here present a novel highly efficient MDS xenotransplantation model, in humanized immunodeficient “MISTRG” mice, expressing humanized M-CSF, IL3/GM-CSF, SIRP alpha, and Thrombopoietin in the Rag^-/-^, IL2R_γ_^-/-^ genetic background from their endogenous murine loci. MISTRG mice have previously been shown to be highly permissive for human hematopoiesis and support robust reconstitution of human lymphoid and myelo-monocytic cellular systems ^14,15^. We demonstrate that primary healthy bone marrow- (BM) and MDS BM-derived CD34+ cells from lower- (IPSS low- and intermediate 1) and higher- (intermediate 2 and high) risk MDS, defined by the number of cytopenias, blast percentage in BM, and cytogenetic abnormalities, efficiently engraft in MISTRG mice and give rise to multilineage hematopoiesis. We demonstrate that MDS patient-derived MISTRG xenotransplants (MDS MISTRG PDX) support the MDS stem cell across all MDS subtypes, replicate the patients’ MDS immunophenotype and histologic features, faithfully reproduce the clonal complexity of the disease at time of diagnosis and along disease progression, and are ideally suited for the testing of targeted therapeutics. Thus, given the high multilineage engraftment efficiency for normal and MDS HSCs and the histologic and clonal fidelity, MISTRG PDX represent a significant advancement over currently available xenotransplantation models and an ideal *in vivo* pre-clinical model for MDS.

## RESULTS

### MISTRG mice engraft healthy adult bone marrow-derived CD34+ hematopoietic stem and progenitor cells

Adult CD34+ hematopoietic stem and progenitor cells (HSPC) engraft with significantly lower efficiency in immunodeficient mice compared to human fetal liver- or cord blood-derived CD34+ cells ^14^. However, the majority of myeloid malignancies and in particular MDS, occur in the aging adult with quantitative and qualitative limitations to the stem cell population of interest. We transplanted healthy BM-derived CD34+ cells from adult patients, in whom BM involvement by their underlying disease was excluded (see **Supplementary Table 1**), intrahepatically into newborn NSG and MISTRG mice irradiated with maximum tolerated doses for each strain (**Fig. 1a**) ^14^. The maximum tolerated radiation in NSG mice is limited due to the inherent DNA repair defect conferred by the *scid* mutation ^16,17^. Samples were CD34 -enriched or CD3 -depleted (**Fig. S1a**), and further purged of mature T-cells by pretreatment with the humanized anti-CD3 antibody OKT3 for prevention of graft versus host disease ^18^. Highest available, rather than a fixed cell number, were injected as equal split- donor grafts into NSG and MISTRG mice, to maximize engraftment for each primary sample.

Analysis consisting of complete blood counts and histology (representative subset), and flow cytometry of peripheral blood (PB), bone marrow (BM), and spleen, was performed at least 12 weeks post-transplantation (with few exceptions were analysis was required sooner). Flow cytometric analysis consisted of assessment of overall human leukocyte engraftment (huCD45) as a function of all (murine and human) leukocytes as well as assessment for human erythroid and megakaryocytic engraftment within the murine and human CD45 negative fraction. "Erythroid lineage” engraftment based on CD45 negativity and high transferrin receptor (huCD71) and glyocphorin A (huCD235) expression was quantitated as %of all single live cells (**Fig. 1b**).

MISTRG mice show significantly higher huCD45+ engraftment in PB and BM than NSG mice (**Fig. 1c, d**) and support enhanced differentiation towards myelopoiesis (**Fig. 1e**) as opposed to lymphopoiesis, modeling a key difference between human and mouse hematopoiesis and as previously shown for mobilized peripheral blood CD34+ HSPCs ^14,19^. CD3+ T-cells represent only a minor fraction within the graft. Interestingly, MISTRG bone marrows show significantly higher numbers of erythroid progenitor cells (CD71^bright^, GPA+) (**Fig. 1f**) as well as megakaryocytes (**Fig.1g**). As previously described, expression of human GM-CSF and M-CSF enhance myeloid maturation with differentiation towards mature granulocytes and macrophages (**Fig.1h, S1b**) with repopulation of bone marrow as well as spleen and non-hematopoietic tissues, such as liver (**Fig. S1c**).

**Figure 1.**
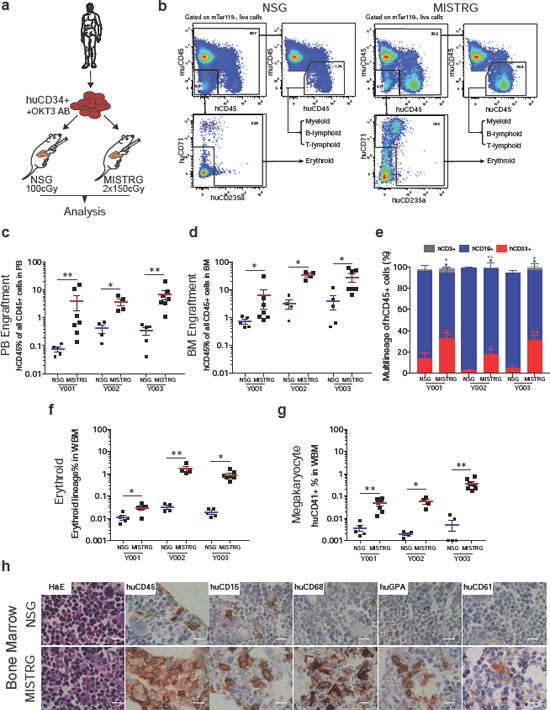
Enhanced engraftment of adult healthy BM derived CD34+ HSPC in human cytokine- knockin MISTRG mice. (**a**) Universal experimental setup. Human BM derived CD34+ HSPCs were pre-incubated with anti-CD3 antibody (OKT3) and injected intrahepatically into newborn (D2-3) NSG or MISTRG mice conditioned with the respective maximum tolerated irradiation doses (NSG 100cGy, MISTRG 2x150cGy). Mice were analyzed 10-17 (healthy BM), 13-30 (MDS), and 9-24 (AML) weeks post transplantation. (**b**) Flow analysis scheme for engrafted NSG or MISTRG mice. Single cell suspensions were stained as in methods and plots gated on live single cells. For analysis of leukocyte engraftment cells were gated on human vs murine CD45 and huCD45 engraftment was calculated as %of all CD45+ cells. For analysis of the erythroid and megakaryocytic lineages the CD45 negative fraction was further analyzed for huCD71^bright/^+ and huCD235+. Erythroid lineage engraftment was calculated as %of total BM cells. (**c, d**) Comparison of overall human engraftment in PB and BM within human grafts in NSG versus MISTRG mice. Individual mice are represented by symbols. (**e**) Relative distribution of myeloid CD33+ (red), B-lymphoid CD19+ (blue), and T-lymphoid CD3+ (Grey) cells as %of human CD45+ cells in NSG vs. MISTRG mice. (**f,g**) Comparison of erythroid and megakaryocytic lineage engraftment in BM of NSG and MISTRG mice. (**h**) BM histology of representative NSG and MISTRG mice from (d). H&E and immunohistochemistry (IHC) stains for huCD45, huCD15, huCD68, huCD235, and huCD61 in NSG (top) and MISTRG BM (bottom row) (scale bars 10μm, original magnification 60x). For detailed sample information see Supplementary Table 1. (In panels c, d, e, f, g data are represented as means ± S.E.M; Mann Whitney test: n.s. not significant, *p<0.05, **p<0.01, ***p<0.001, ****p<0.0001).

In summary, MISTRG mice support superior healthy adult BM xenografts with tri-lineage representation.

### MISTRG mice efficiently support PDX from low- and high-risk MDS with multi-lineage output

NSG mice have represented a major breakthrough in xenotransplantation studies due to the lack of mature murine T, B, and functional NK cells ^20^ and the presence of the *Sirpα* gene polymorphism, allowing enhanced binding of the mSirpα to human CD47 ^21,22^. However, engraftment of MDS BM-derived CD34+ HSPCs remains a challenge, despite several alterations to NSG mice and the transplantation protocol ^4-7,9-12^. We engrafted MDS CD34+ (or CD3-depleted) BM cells into NSG and MISTRG recipients as split donor grafts, as in Fig. 1a. To avoid a priori exclusion of lower-risk MDS samples or patient samples with low cell numbers, CD34+ cells injections for different samples ranged from 0.5x10^5^ to 1x10^6^ cells per recipient mouse, while maintaining the same cell number for all recipients within each experiment (for detailed patient and sample information see **Supplementary Table 1**).

When plotting engraftment in all mice against injected CD34+ cell number, it is evident, that a minimum number of 1x10^5^ CD34+ cells / mouse was required for a reliable engraftment (**Fig. 2a**). Interestingly, increasing cell numbers resulted in improved engraftment in MISTRG while engraftment in NSG recipients remained limited. Although all recipients engrafted above 0.01%, the minimum engraftment threshold set in several studies, for the purpose of pre-clinical modeling a higher engraftment threshold may prove advantageous. When comparing all split donor graphs, engraftment > 1%was achieved in 85%of MISTRG and in 52%of NSG mice. Importantly, engraftment levels >10%, more likely to reliably afford preclinical studies, were achieved in 53%of MISTRG but in less than 10%of NSG mice (**Fig. 2b**). MISTRG consistently resulted in higher engraftment than NSG for all MDS subtypes in peripheral blood (top row) and bone marrow (bottom row) (**Fig. 2c - e**). Only 2 out of 28 samples (MDS-MLD Y006, MDS-EB2 Y018), injected at <1x10^5^CD34^+^ cells/mouse displayed BM engraftment levels < 1%in MISTRG. CD3-depletion of primary MDS BM samples, similar to CD34-enrichment, resulted in similar engraftment levels in PB and BM (**Fig. S2a, b**) and myeloid predominant grafts in MISTRG mice (**Fig. S2c**).

**Figure 2.**
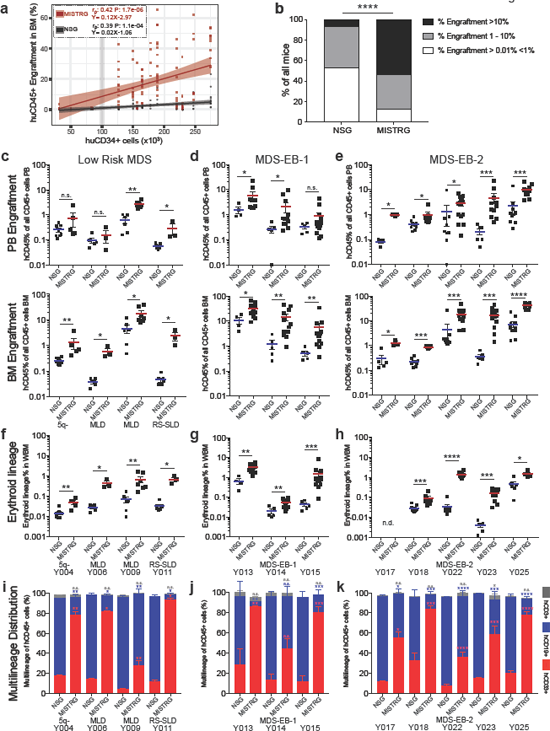
Enhanced engraftment of higher- and lower-risk MDS in MISTRG mice. (**a**) Split donor huCD45+ BM engraftment in NSG (black) versus MISTRG (red) mice plotted against CD34+ cell number injected/mouse. Individual mice are represented by symbols. Linear regression,Pearson correlations and p-values of %engraftment to CD34+ cell number in NSG (r=0.39, P<0.0001) vs MISTRG (r=0.42, P<0.0001) are displayed. (**b**) Percentage of transplanted mice with huCD45+ BM engraftment levels > 0.01%<1%, 1-10%and >10%for split-donor grafts in NSG (59/111,44/111 and 8/111, respectively) and MISTRG (20/154, 51/154 and 83/154, respectively) mice (Fisher's exact test, ****p<0.0001 for NSG vs MISTRG). (**c-k**) Analysis of huCD45 and lineage engraftment was performed as detailed in Fig. 1b > 13 weeks post transplantation. **(c, f, i)** Analysis of MDS - 5q, -SLD, -MLD, and -MLD-RS - engrafted NSG and MISTRG mice. (**d, g, j**) Analysis of MDS-EB-1 - engrafted NSG and MISTRG mice. (**e, h, k**) Analysis of MDS-EB-2 - engrafted NSG and MISTRG mice. MISTRG afford significantly higher engraftment than NSG in high- and low-grade MDS. (**c-e**) Comparison of overall human CD45+ leukocyte engraftment in PB (top) and BM (bottom) in NSG versus MISTRG mice. **(f-h)** Comparison of erythroid lineage engraftment in BM in NSG and MISTRG mice. Individual mice are represented by symbols with means ± S.E.M. (**i-k**) Relative distribution of myeloid CD33+ (red), B-lymphoid CD19+ (blue), and T-lymphoid CD3+ (Grey) cells as %of human CD45+ cells in NSG vs. MISTRG mice. Stacked bar graphs represent means ± S.E.M. Mann Whitney test; n.s. not significant, *p<0.05, **p<0.01, ***p<0.001, ****p<0.0001. For detailed patient sample information see Supplementary Table 1.

Importantly, we here show for the first-time engraftment of primary adult MDS-derived erythropoiesis (defined by huCD71^bright^ and huCD235 positivity among CD45^neg^ cells) and megakaryopoiesis (huCD41+) in immunodeficient mice, with significantly higher representation in MISTRG mice for all subtypes of MDS (**Fig. 2f-h and Fig. S2d-f**). As described previously for normal hematopoiesis, CD34+ cells from MDS bone marrow give rise to myeloid-predominant, while NSG mice give rise to lymphoid-predominant grafts (**Fig. 2i-k**). Expression of human M-CSF, GM-CSF, and IL-3 further enhances maturation of MDS-derived myeloid cells with differentiation profiles close to the patients’ phenotypes (**Fig. S2g**). To assure that xenografts are derived from the malignant MDS clone we performed mutational analysis by targeted exome sequencing of patient samples and corresponding murine cell-depleted patient-derived xenografts. Presence of corresponding driver mutations at equivalent variant allele frequencies (VAF) confirmed engraftment of MDS-derived hematopoiesis (**Supplementary Table 2**).

The engraftment persisted until the time of analysis, > 12 weeks post-transplantation, without compromising development of anemia or thrombocytopenia in recipients mice (data not shown) or differences in survival between MISTRG and NSG mice (**Fig S2h**), with similar engraftment in female and male mice of the respective strains (**Fig. S2i**). Analysis of spleen and non-hematopoietic tissues (shown for liver) confirms functional differences between healthy BM, MDS, and AML xenografts. While healthy BM xenografts engraft BM, liver, and spleen and give rise to resident tissue macrophages in in all three tissues (**Fig. S2j and S3a**), in age-matched patient-derived MDS xenografts these populations are mostly absent from the spleen and the liver, consistent with a functional defect of the myeloid lineages in MDS (**Fig.S2j and S3b**). This is in stark contrast to myeloid leukemia where immature blasts infiltrate spleen and liver (**Fig. S2j and S3c**).

In summary, MISTRG mice support superior long-term, multi-lineage engraftment of MDS cytokine- humanization enhances myeloid lineage maturation and supports erythroid and megakaryocytic lineage development. Furthermore, MISTRG PDX replicate functional differences between healthy, MDS, and leukemia-derived hematopoiesis.

### The MISTRG humanized stem cell niche allows propagation of MDS HSC via serial transplantation

HSCs are critically dependent on the stem cell niche. MDS HSC are dysfunctional and their *in vitro* and *in vivo* propagation has been elusive to date. We hypothesized that cytokine humanization of the HSC niche would afford homing and engraftment of primary patient-derived MDS long-term HSC in MISTRG mice capable of serial repopulation. Human thrombopoietin is essential for stem cell function ^23,24^. IL3, GM- or M-CSF are not directly implicated in stem cell maintenance, but via their role in myeloid cell development, such as BM macrophages, they may indirectly supply additional niche signals ^25,26^. We assessed human versus murine cytokine expression (**Fig. 3a, b**) in MDS (**Fig. S4a**) and murine MISTRG and NSG (**Fig. S4b**) BM-derived mesenchymal stromal cell (MSC) cultures. Human and MISTRG MSC, but not NSG MSC express human THPO, GM-CSF, and M-CSF instead of their murine counterparts. IL3, as expected, is not expressed in MSC (data not shown)^27^.

**Figure 3.**
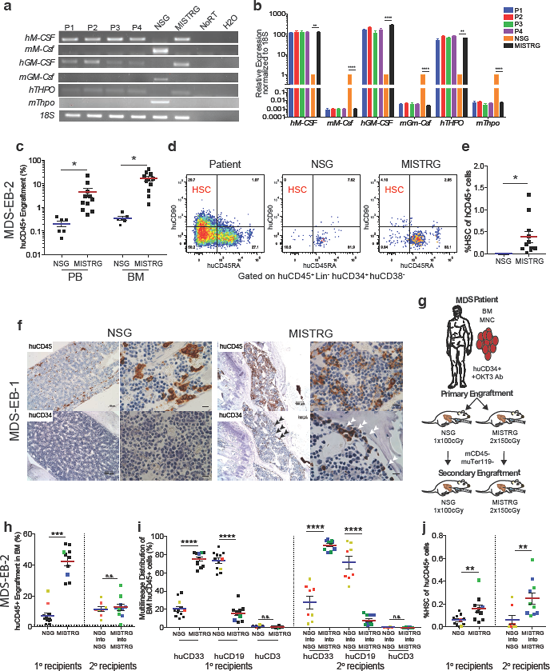
MISTRG support phenotypic and functional MDS stem cells with long-term multilineage engraftment potential. (**a-b**) MISTRG BM-derived mesenchymal stromal cells (MSCs) express human in place of murine cytokines. RT-PCR (**a**) and Q-RT-PCR (**b**) detection of human versus murine cytokine expression in 4 patient-derived (P1-4) and representative NSG and MISTRG MSC cultures. (Representative experiment of n=3 independent experiments, statistics represent One way ANOVA with Tukey’s multiple comparison calculations; **p<0.01, **** P<0.0001). (**c-f**) MISTRG engraft phenotypic MDS stem cells. (**c**) Overall human cell engraftment and (**d**) representative FACS plots and (**e**) quantification of HSC representation (lin^-^CD34+CD38^-^CD45RA^-^CD90+ of huCD45+) of corresponding patient (high-risk MDS-EB2, Y023), and NSG and MISTRG xenografts. (**f**) Representative IHC for huCD45 and huCD34 distribution in NSG (of n=5) and MISTRG (of n=12) BM engrafted with MDS-EB-1 (Y014; scale bars for low-power field: 100μm, original magnification 10X; high-power field: 10μm, original magnification 60X). (**g-j**) MISTRG engraft functional MDS stem cells. (**g**) Secondary xenotransplantation experimental setup. (**h-j**) Primary and secondary transplantation of high-risk MDS-EB2 (Y025) comparing (**h**) overall engraftment in BM and (**i**) multi-lineage representation in BM of primary and secondary NSG and MISTRG recipients. (**j**) Comparison of phenotypic HSC %in NSG vs. MISTRG primary and secondary recipients as in (d, e). Individual mice are represented by symbols with means ± S.E.M.; symbols for corresponding 1° and 2° recipient mice are color-coded; statistics represent Mann-Whitney test; n.s. not significant, *p<0.05, **p<0.01,***p<0.001, ****p<0.0001. For patient information see Supplementary Table 1.

We next determined whether MISTRG mice engraft human HSC via phenotypic and functional assays. In addition to the overall increased engraftment (**Fig. 2, 3c, S4c)**, MISTRG support phenotypic HSC as evident by flow-cytometric analysis (**Fig. 3d, e**). CD34+ cells localize along the trabecular bone (**Fig. 3f and S4d**). The clonality of each MDS graft was verified by targeted exome sequencing in representative mice (**Fig. S4e, f and Supplementary Table 2**).

Phenotypic identification of HSCs is insufficient to prove stem cell engraftment. Long-term engraftment (> ~16 weeks) and functional assessment in the form of secondary engraftment are critical. Previous studies have shown successful secondary transplantation of AML ^28,29^ and more recently CMML/JMML in NSG and NSG- SGM3 mice ^30^ but no study has shown successful serial transplantation of MDS. We therefore first tested secondary transplantation of a higher-risk MDS sample according to our standard protocol (**Fig. 3g**). NSG and MISTRG mice were engrafted with high-risk MDS-EB-2 with complex karyotype (Y025) and maintained for > 16 weeks. At time of analysis BM from the highest engrafting mice was enriched for human cells via bead depletion of murine CD45+ and Ter119+ cells (**Fig. S4g, h**) and transplanted intrahepatically into irradiated newborn mice. Secondary recipient mice were analyzed >12 weeks post 2° transplant. Primary MISTRG recipient mice again showed higher overall engraftment levels (**Fig. 3h**) with myeloid predominance (**Fig. 3i**). Cytogenetic analysis of sorted human cells from NSG and MISTRG primary recipients showed the same complex cytogenetic abnormalities as the patient’s (**Fig. S4i**). Interestingly, secondary NSG and MISTRG mice showed similar huCD45+ engraftment in bone marrow (**Fig. 3h**), but again myeloid predominance prevailed in MISTRG, while lymphoid predominance prevailed in NSG secondary recipients (**Fig. 3i**). Notably, MISTRG xenografts contained significantly higher HSC than NSG mice (**Fig. 3j, S4j**) suggestive of a favorable stem cell niche environment.

We next compared the ability of MISTRG mice to propagate karyotypically normal (NK) intermediate risk MDS (Y022, MDS-EB-2, NK, IPSS int-2, **Supplementary Table 1**). MISTRG 1° recipients served as donors for NSG and MISTRG 2° recipients (**Fig. 4a**). Both strains’ 2° recipients successfully propagated clonal MDS for > 16 weeks from 1° MISTRG donors (15 weeks), however, 2° NSG engraftment levels were < 1% (**Fig. 4a**) with significantly lower percentages of HSC (**Fig. 4b**) and favorable myeloid representation in MISTRG 2°recipients (**Fig. 4c**). Targeted exome sequencing of primary MISTRG and secondary MISTRG and NSG recipients verified propagation of the patient’s clonal MDS (**Fig. 4d, Supplementary Table 2**). Given the similar VAFs between MISTRG and NSG mice suggests that B-cells in MDS xenografts are derived from the MDS clone. Similar results were obtained with serial transplantation of lower-risk MDS. NSG and MISTRG mice were engrafted with intermediate-1 risk MDS (Y013, MDS-EB1, NK, IPSS int-1, **Supplementary Table 1**) and their murine-cell depleted bone marrow engrafted after 22 weeks into irradiated recipients of the respective genotypes. Again, NSG secondary recipients showed low-level engraftment (~1%), while MISTRG-derived cells successfully repopulated numerous MISTRG secondary recipients at significantly higher engraftment levels (p<0.001; **Fig. 4e**). These engraftment outcomes are again reflected in the phenotypic stem cell frequency in 1° and 2° NSG and MISTRG grafts with significantly higher percentage of HSC in MISTRG mice (p< 0.05**; Fig. 4f**) with myeloid predominant output **Fig. 4g**) MDS clonality of primary and secondary grafts in MISTRG and NSG recipients was again confirmed by targeted exome sequencing (**Fig. 4h**).

**Figure 4.**
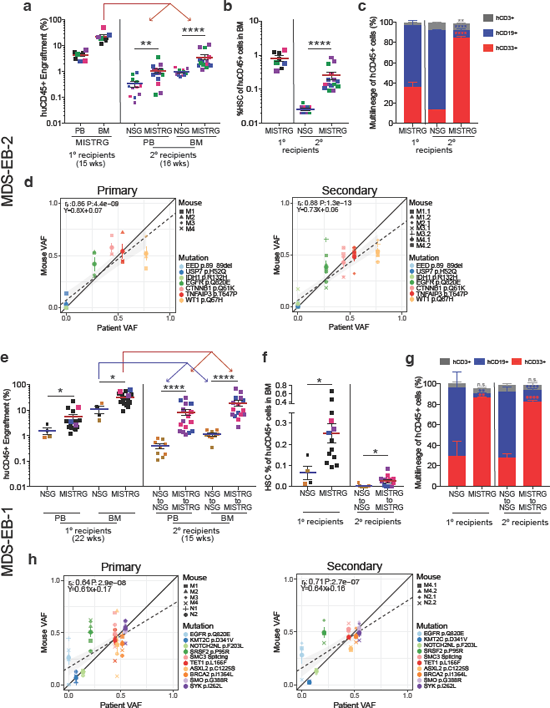
MISTRG faithfully propagate clonal long-term HSC. Serial engraftment, lineage output, and clonal representation of (**a-d**) IPSS intermediate-2 MDS-EB-2 (Y022) and (**e-h**) IPSS intermediate-1 MDS-EB1 (Y013) engrafted NSG and MISTRG mice. (**a**) Human CD45+ engraftment in MISTRG PB and BM and comparison of engraftment in secondary NSG and MISTRG recipients. (**b**) Phenotypic HSC (Lin^-^ CD38^-^ CD34+ CD45RA^-^ CD90+ %of huCD45+) of primary MISTRG and secondary NSG and MISTRG recipient mice (Individual mice are represented by symbols with mean ± S.E.M.. Corresponding primary and secondary recipient mice are color-coded. Mann Whitney test with *p<0.05, **p<0.01, ***p<0.001, and ****p<0.0001 for aggregate NSG vs. MISTRG). (**c**) Relative distribution of myeloid CD33+ (red), B-lymphoid CD19+ (blue), and T-lymphoid CD3+ (Grey) cells as %of human CD45+ cells in 1° and 2° NSG vs. MISTRG recipient mice. Stacked bar graphs represent means ± S.E.M. Mann Whitney test; n.s. not significant, *p<0.05, **p<0.01, ***p<0.001, ****p<0.0001, for aggregate NSG vs. MISTRG. (**d**) Clonality was determined in representative primary and secondary MISTRG recipients with engraftment levels > 1%via targeted exome sequencing. Variant allele frequencies (VAFs) in primary and secondary recipients were plotted against the corresponding patient’s. Individual mice are represented by symbol shape and mutations are color-coded. Linear regression, Pearson correlations and p-values between patient and xenograft VAF are displayed. (**e**) Comparison of human CD45+ engraftment in PB and BM in primary NSG and MISTRG mice and secondary recipients of mice from the respective primary strain. (**f**) Phenotypic HSC in primary and secondary recipient mice. Statistic were calculated as in (a, b). (**g**) Relative distribution of myeloid CD33+ (red), B-lymphoid CD19+ (blue), and T-lymphoid CD3+ (Grey) cells as %of human CD45+ cells in NSG vs. MISTRG mice. Statistic were calculated as in (c). (**h**) VAFs in primary and secondary NSG and MISTRG mice engrafted > 1%. Individual NSG and MISTRG mice are represented by symbol shape and mutations are color-coded. Statistic were calculated as in (d). For detailed patient information see Supplementary Table 1.

Overall, these data suggest, that MISTRG not only provide overall superior engraftment in primary recipients with propagation of the MDS stem cell, but also as secondary recipients. This may at last fill the unmet need for MDS PDX for the study of MDS, allowing the development and testing of novel therapies.

### MISTRG mice replicate MDS heterogeneity, myeloid dysplasia, and clonal evolution

Although several murine models of MDS have been generated, the finding of dysplasia is rare and frequently subtle (reviewed in ^31^). Currently available xenotransplantation models have not been shown to replicate myelodysplasia, the essential diagnostic criterion for MDS, nor to support development of erythro- and megakaryopoiesis, two of the three principal cell lines affected in MDS ^32,33^. We engrafted two MDS samples with prominent dysplasia of the megakaryocytic and erythroid lineages, respectively, to evaluate dysplasia in human MDS xenografts. MISTRG mice engrafted with sample Y019 (MDS-EB-1 with normal karyotype, **Fig. 5a,** left) displayed numerous dysplastic megakaryocytes and reticulin fibrosis, faithfully replicating the patient’s MDS dysplastic features (**Fig. 5b**). MISTRG mice engrafted with the same patient’s sAML sample obtained at time of disease progression (Fig. 5a; Y028, sAML, NK) did not show these features (data not shown). Targeted exome sequencing of the MDS xenografts confirmed derivation from the patient’s DNMT3a-mutant MDS clone (**Fig. 5c**). Interestingly, an isocitrate dehydrogenase 1 (IDH1) mutation was identified in several of the MISTRG mice (VAF 18-32%) engrafted with the patient’s initial MDS diagnosis sample (MDS-EB-1, **Fig. 5c**). This IDH1 mutation was initially not identified in the patient at the time of MDS diagnosis, but present at the time of disease progression to sAML (sAML, VAF 24%, **Fig. 5c**). Re-sequencing detected the IDH1 R132C mutation in the MDS diagnosis sample at a VAF of ~1% (**Fig. 5c,** middle panel, **Supplementary Tables 1 and 2**). Interestingly, in the sAML engrafted MISTRG mice a new NRAS G12S mutation defined the dominant clone, again detectable in the patient’s sAML at a VAF <5%upon resequencing, in addition to the dominant DNMT3a and IDH1 mutations. RAS mutations have been described as a potential mechanism of resistance to mutant IDH inhibitor treatment ^34^ and identification of these mutant clones in a pre-clinical MDS PDX may thus guide the use of pre-emptive combination regimens.

**Figure 5.**
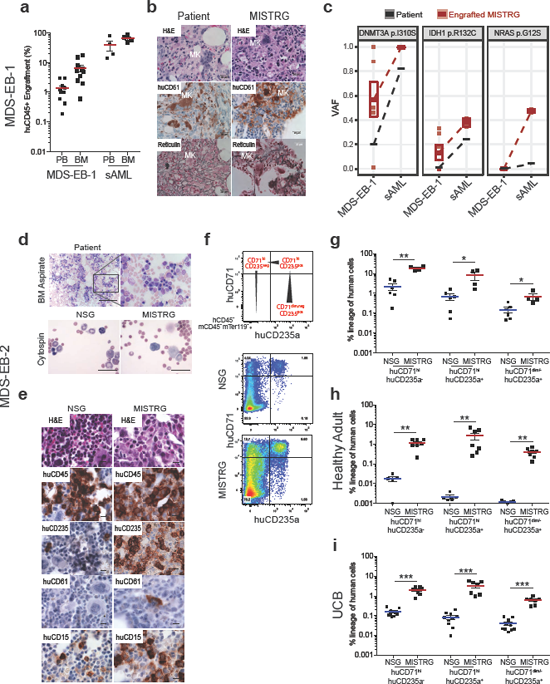
MISTRG replicate myelodysplasia and clonal evolution upon disease progression. (**ac**) MISTRG engrafted with consecutive MDS-EB-1 and sAML samples from the same patient (Y019 and Y028, respectively). (**a**) Overall (huCD45+) engraftment in PB and BM. Individual mice are represented by symbols, with means ± S.E.M. (**b**) Histology from MDS-EB-1 (Y019) diagnostic BM and representative engrafted MISTRG BM. H&E and huCD61 stains reveal human megakaryocytic dysplasia and reticulin stain reveals bone marrow fibrosis (high power magnification scale bars 20μm). (**c**) Targeted exome sequencing results from MISTRG xenografted with same patient’s primary MDS-EB-1 diagnosis samples and sAML at time of disease progression. For each mutation, variant allele frequencies (VAFs) are shown for the patient (black) and representative MISTRG (Red) mice with engraftment levels >1%. Mean VAF values between MDS-EB-1 and sAML are connected by lines. (**d-f**) MISTRG xenografted with MDS-EB-2 (Y025) BM with inverted Myeloid: Erythroid ratio and marked erythroid dysplasia. (**d**) Patient BM aspirate (top) and cytospins from engrafted NSG and MISTRG BM after murine CD45+ and Ter119+ depletion (bottom); (for overall engraftment see Fig. 2 e, h, k, Y025). (**e**) Representative BM histology from representative NSG and MISTRG recipients engrafted > 1%. (**f**) Representative FACS plots of erythroid lineage differentiation based on huCD71 and huCD235 expression in huCD45^-^ muCD45^-^ mTer119^-^ cells (huCD71^hi^huCD235a^-^ (ProErythroblasts (EB)), huCD71^hi^huCD235a+ (Basophilic EB/ Normoblasts), huCD71^-^huCD235a+ (reticulocytes, RBC)). Comparison of MDS-EB2 (Y025) (**g**), healthy adult (Y003) (**h**), and umbilical cord blood (UCB) (**i**) erythroid lineage differentiation in NSG and MISTRG recipient mice. (For overall engraftment see Fig. 1c and Fig. S5a.) Individual mice are represented by symbols with mean ± S.E.M.; statistics represent Mann Whitney test; *p<0.05, **p<0.01, ***p<0.001, for aggregate NSG vs. MISTRG.

The second patient’s BM aspirate (MDS-EB-2, Y025) revealed significant erythroid hyperplasia with erythroid dysplasia and maturation defect (**Fig. 5d** top). Bone marrow aspirate/cytospins of engrafted primary MISTRG and to a much lesser extent of engrafted NSG revealed erythroid precursors with signs of dysplasia, such as binuclear forms (**Fig. 5d** bottom). Importantly, BM histology revealed prominent development of huCD235+ erythroid progenitors in MISTRG mice (**Fig. 5e**), confirmed by flow cytometric determination of huCD71 ^pos^ and huCD235^pos^ (gated on CD45^neg^, mTer119^neg^ and huCD45^neg^ cells) erythroid development as shown in Fig. 5f and quantitated in **Fig. 5g**. Interestingly, compared to xenografts from healthy BM- (Fig. 5h) and human umbilical cord blood-derived CD34+ HSCPs (Fig. 5i), erythroid differentiation is left-shifted (to huCD71^hi^ huCD235^-^) in MDS-xenografts with enhanced maturation in healthy BM and UCB (**Fig. 5h,i**). These differences are not evident in NSG mice given the significantly lower representation of the erythroid lineage, despite successful engraftment of the respective samples (MDS Y025, Fig. 2e, h, k; Healthy BM, Y003, Fig. 1c-g, UCB **Fig. S5a-d**;). Importantly, erythroid lineage differentiation is always significantly higher in MISTRG than NSG mice, and present in secondary recipients (**Fig. S5e-i**), confirming that it is derived from the MDS stem cell.

In summary, we here show that MISTRG MDS PDX may predict clonal evolution upon disease progression with faithful modeling of the diagnostic dysplasia. Importantly, we present the first MDS PDX model that affords the study of MDS erythropoiesis and megakaryopoiesis *in vivo.*

### MISTRG MDS PDX allow preclinical modeling of targeted therapeutics

Targeted therapeutics provide novel opportunities for the treatment of MDS, but to date have failed to cure MDS. Recently, inhibitors of mutant IDH1/2 have entered clinical trials, and early data suggest that they result in blast differentiation, but fail to abrogate the mutant clone in the majority of patients ^34-36^. While transgenic murine models can provide proof of principle data, patient-derived xenografts are critical to evaluate efficacy against complex clonal hematopoietic malignancies such as MDS, and are likely to hasten development of valuable combination therapies.

We transplanted MISTRG mice with IDH2^R140Q^ -mutant MDS-EB-2 CD34+ cells (Y021, **Supplementary Table 1**) and treated engrafted mice with either vehicle or enasidenib via oral gavage for 30 days. Mice were assigned to enasidenib or vehicle 16 weeks post-transplantation based on equal engraftment levels as determined by BM aspiration (data not shown). Activity of enasidenib was verified *in vitro* via measurement of 2-hydroxy-glutarate (2-HG) levels in IDH2-wildtype (WT) and -mutant (MUT) expressing human erythroid leukemia (HEL) cell lines (**Fig. S6a**) and primary AML (**Fig. S6b,** Y028, Y031). Enasidenib efficiently suppressed 2-HG production and inhibited proliferation of IDH2^R140Q^- and IDH2^R172K^-mutant but not IDH2- wildtype AML cell lines (data not shown) and IDH2^R140Q^-mutant primary AML compared to vehicle and WT AML (**Fig. S6c**). Enasidenib treatment resulted in differentiation of IDH2 mutant myeloid blasts (**Fig. S6d**).

Treatment with enasidenib, but not vehicle, resulted in myeloid differentiation in the IDH2^R140Q^ MDS-EB-2 - engrafted MISTRG mice (**Fig. 6a**, hCD68 and hCD15 and Fig. 6b). Overall engraftment levels were significantly reduced in enasidenib treated animals when compared to pre-treatment and vehicle-treated mice (**Fig. 6c**). Of note, enasidenib treated mice also exhibited increased numbers of CD41^+^ platelets in PB and dysplastic megakaryocytes in BM compared to vehicle treated mice (**Fig. 6a** (huCD61), **Fig. 6d**). Plasma 2-HG levels *in vivo,* elevated pre-treatment and in vehicle treated MISTRG mice, were significantly suppressed after administration of enasidenib (**Fig. 6e**). Variant allele frequencies of mutations identified in the patient were represented in all MISTRG mice and not significantly altered by enasidenib treatment (**Fig. 6f, Supplementary Table 2**).

**Figure 6.**
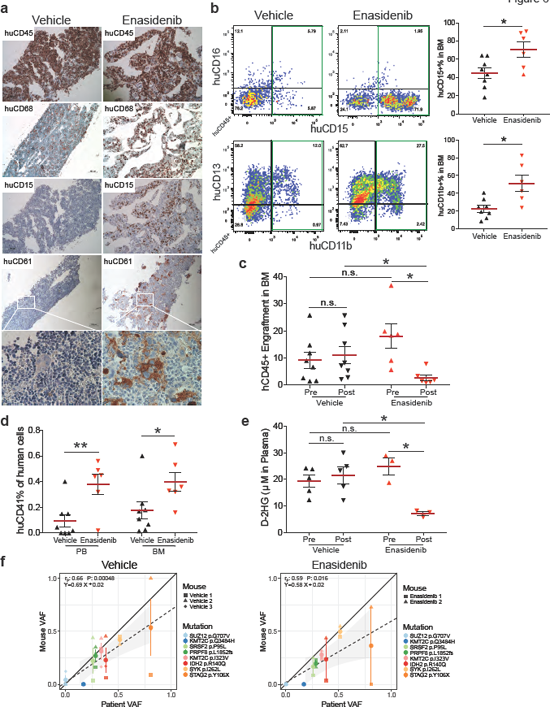
MISTRG replicate granulocytic and megakaryocytic differentiation in response to inhibition of mutant IDH2. *In vivo* treatment of mutant IDH2 R140Q in MDS-EB-2 (Y021) engrafted MISTRG mice with the IDH2^MUT^ inhibitor enasidenib. (**a**) Representative histologic images of vehicle (n=8, left) and enasidenib (n=6, right) treated mice engrafted with MDS-EB-2 (Y021). IHC stains for huCD45, huCD68, huCD15, and huCD61 (scale bars 10μm, original magnification 60X). (**b**) Representative FACS plots showing myeloid maturation in response to enasidenib and quantitation of huCD15+ and huCD11b+ expression in vehicle versus enasidenib treated MISTRG mice. (**c**) Comparison of human engraftment in BM from vehicle (n=8) and enasidenib (n=6) treated MISTRG mice. (**d**) Quantitation of huCD41+ expression in PB and BM from vehicle (n=8) and enasidenib (n=6) treated MISTRG mice (**e**) Quantitation of D-2HG in plasma of pre- and post-administration of Vehicle or Enasidenib. Individual mice are represented by symbols with mean ± S.E.M.; statistics represent Mann Whitney test; n.s. not significant, *p<0.05, **p<0.01 for aggregate NSG vs. MISTRG. (**f**) Representation of VAFs of driver mutations in vehicle (left) or enasidenib (right) treated MISTRG (y-axis) plotted against the patient’s VAFs (x-axis). Individual mice represented by symbol shape, mutations color coded. Linear regressors, pearson correlations and p-values between patient and xenograft VAF are displayed.

MISTRG PDX represent the first MDS pre-clinical model that allows to test not only for cytotoxic but also for differentiating effects of targeted therapeutics and capturing multi-faceted responses relevant to clinical success.

## DISCUSSION

MDS is a disease of the hematopoietic stem cell and studies of MDS have been hampered by the inability to expand HSCs in general and MDS stem cells in particular. There is an unmet need for an *in vivo* pre-clinical model to accelerate development of novel treatments for a disease where allogeneic stem cell transplantation currently represents the only cure. Mouse models only partially recapitulate the genetic and epigenetic complexity of patients’ MDS. Prior xenotransplantation models have been hampered by preferential engraftment of the remnant normal hematopoiesis ^7^, transient engraftment, ^6^, and low efficiency ^37^. Cytokine- humanization via transgenic expression in the NSG-SGM3 mice, while advantageous in AML ^9^ and other myeloid malignancies ^30^, impairs stem cell function ^8,11^ and provides limited advantages over NSG mice for MDS engraftment ^37^. Co-injection of human MSC may provide transient support to MDS HSC ^12,37^ and generation of a human niche via growth of human MSC-derived ossicles may afford improved engraftment of HSCs ^13,38^ and difficult-to-engraft leukemias ^38^, but applicability in pre-clinical models at a large scale is likely limited due to technical complexity.

MISTRG mice were engineered to express key non-crossreactive human cytokines from the endogenous murine loci in place of their murine counterparts, thereby providing temporally and spatially physiologic expression of human cytokines. In addition, lack of murine cytokines reasonably provides additional benefit to human hematopoiesis by rendering the stem and progenitor niches in the BM less hospitable to murine HSPC. This is likely to particularly critical to adult HSCs that have markedly lower proliferative and self-renewal capacity than their fetal liver and UCB counterparts (reviewed in ^39^). In addition, MDS stem cells frequently fail to give rise to colony forming units (CFU) *in vitro,* a manifestation of their defective proliferative and differentiation capacity. Research material from human BMs is limited and worse in aging marrows that are characterized by progressively lower cellularity.

Here, we present for the first time a highly efficient and versatile xenotransplantation model for MDS. We show that MISTRG mice can be engrafted with as few as ~1.5x10^5^ MDS BM-derived HSPC. Higher engraftment levels clearly improve MISTRG utility as a model. Over 80%of MISTRG mice engraft when a threshold of 1%BM huCD45+ cells is set. More remarkably, over 50%of MISTRG mice, compared to fewer than 10%of NSG mice, engraft above a threshold of 10%BM huCD45+ cells. In addition, MISTRG mice persistently show improved myeloid representation and differentiation, as evident by both flow-cytometry and histologic evaluation.

Study of adult erythropoiesis and megakaryopoiesis in *in vivo* models has been elusive to date. MDS is characterized by cytopenias in the peripheral blood, left-shifted myeloid maturation, and erythroid and megakaryocytic dysplasia ^3^. Very little is known about the causes of the phenotypic heterogeneity in MDS and genotype-phenotype studies would greatly advance our mechanistic understanding of this complex entity. To date, immunodeficient mouse models have supported erythro- and megakaryopoiesis solely from fetal liver- and cord blood-derived HSPCs ^40,41^, further enhanced by mutation of the murine ckit receptor conferring impaired function to murine stem cells ^42-44^ and likely murine erythropoietic progenitors ^45^. None of these models have supported erythro- and megakaryopoiesis from adult HSPCs. We propose that while cytokine humanization directly enhances overall engraftment and myeloid maturation, lack of the corresponding murine cytokines impairs murine hematopoiesis, thereby synergistically promoting human HSPC competitiveness in the mouse niche. As a result, we here show for the first time development of both erythropoiesis and megakaryopoiesis from healthy adult and MDS BM in a murine host. Intriguingly, MDS - engrafted MISTRG mice replicate the patient’s megakaryocytic dysplasia, reticulin fibrosis, and erythroid dysplasia, making MISTRG MDS PDX uniquely suited to assess MDS-associated abnormalities in all three myeloid lineages.

We have previously reported that MISTRG life-span is limited in fetal liver engraftment due to destruction of murine RBC and platelets by human macrophages ^14^. Interestingly, MDS-engrafted MISTRG lack significant development of anemia (data now shown) and their life-span is similar to that of engrafted NSG mice (**Supplementary Table 1**). One possible explanation is the lower engraftment compared to fetal liver HSPC. Nevertheless, mice engrafted with normal adult CD34+ cells with similar engraftment levels to MDS engrafted mice show evidence of hemophagocytosis by human macrophages (^19^ and data not shown). In contrast to normal CD34+ engrafted MISTRG, MDS PDX lack human derived tissue macrophages in spleen and liver, confirming a functional defect of MDS-derived mature myeloid cells that likely also lack *in vivo* hemophagocytic activity. Interestingly, MDS blasts, unlike in AML, do not infiltrate non-hematopoietic tissues, functionally distinguishing MDS also from AML in MISTRG PDX.

Cytokine humanization does not alter the lack of mature human red blood cells (RBC) and the low human platelet percentage in the peripheral blood as also shown previously ^14,19^. Administration of human erythropoietin has shown no benefit in this regard ^46^ as the defect lies in RBC and platelet destruction by the murine innate immune system. Thus modulation of the murine innate immune system will be necessary to promote mature human cell persistence in peripheral blood, transiently achieved by administration of liposome-encapsulated clodronate ^41,47^.

MDS is a clonal hematopoietic stem cell disorder and reliable engraftment of the malignant HSC is essential for high qualitypre-clinical studies of disease biology and response to therapeutics. While phenotypic evidence of HSC can suggest their presence, functional stem cell assays are critical. Serial transplantation represents the gold-standard functional hematopoietic stem cell assay. We here show successful serial transplantation of MDS into MISTRG secondary recipients with faithful representation of the clonal composition, lineage representation, and dysplastic features of the parental patients’ BM in primary and secondary recipients. Importantly, MISTRG mice allow the expansion of xenografts from one primary into several secondary recipients, essential for pre-clinical modeling and therapeutic testing.

We interrogated the utility of the MISTRG MDS PDX model in the testing of targeted therapeutics, specifically inhibition of mutant IDH2. Early clinical studies have shown that enasidenib, an oral inhibitor of mutant IDH2, results in differentiation of mutant myeloblasts without abrogation of the mutant clone in the majority of patients. We here show for the first time, in an *in vivo* MDS PDX model, differentiation towards dysplastic megakaryocytes and myeloid maturation with preservation of the clonal composition of the graft. The MISTRG MDS PDX model is ideally suited for the systematic study of targeted therapeutics alone and in combination with other agents. Concurrent targeted exome sequencing may allow to predict ideal combination regimens for individual patients.

In summary, we here present a highly efficient, faithful MDS PDX model, ideally suited for the study of MDS biology, the development of novel treatment approaches, and adaptation of patient-specific regimens in the era of precision medicine.

## MATERIALS and METHODS

### Human Progenitor Cell Isolation

Peripheral blood, BM, and umbilical cord blood were obtained with donor’s written consent. All human studies were approved by the Yale University Human Investigation Committee and by the West Haven Veterans Affairs Human Investigation Committee.

Human BM, cord-blood, and peripheral blood samples were ficolled (GE Healthcare, Munich, Germany) and mononuclear cells cryopreserved within 24 hours after collection in FBS/10%DMSO. Samples were CD34 - enriched with the CD34-Microbead-Kit or T cell - depleted via negative selection with the CD3-Microbead-Kit (Miltenyi-Biotech, Bergisch-Gladbach, Germany). CD34 - enriched or CD3 - depleted HSPCs were incubated with a murine anti-human CD3 antibody (clone Okt3, BioXCell, NH, USA) at 5μg / 100μl for 10 minutes at RT prior to injection into mice.

### Generation and Analysis of MISTRG PDXs

*Mouse breeding and xenografting:* MIS^h/h^TRG mice with homozygous knockin-replacement of the endogenous mouse *Csf1, Il3, Csf2, Tpo,* and *Sirpa* with their human counterparts were bred to MITRG mice to generate human cytokine homozygous and *hSIRPA* heterozygous mice ^14,15^. NSG mice were obtained from Jackson laboratory. MIS^h/m^TRG (labeled MISTRG throughout the manuscript) and NSG mice were maintained on continuous treatment with enrofloxacin in the drinking water (0.27 mg/mL, Baytril, Bayer Healthcare). Newborn MISTRG or NSG mice (1 to 3 days of age) were sublethally irradiated (X-ray irradiation with X-RAD 320 irradiator; MISTRG 2 x 150cGy 4 hours apart, NSG 1 x 100cGy). Equal numbers of split donor MDS BM CD34- selected or CD3-depleted (as indicated) were injected intrahepatically in a volume of 20ul into with a 22-gauge Hamilton needle (Hamilton, Reno, NV). Mice were analyzed at least 12 weeks post transplantation and only sooner if moribund. For secondary transplantation, human cells were isolated from primary recipient BMs and depleted of murine cells via negative selection of murine CD45+ and Ter119+ cells by magnetic labeling with biotin-anti-muCD45 (clone 30-F11, Biolegend, San Diego, CA) and muTer119 (clone TER-119, Biolegend) and BD IMag Streptavidin Particles (BD Biosciences, San Jose, CA).

*Flowcytometric Analysis:* Engraftment of human CD45+ cells and their stem cell, progenitor, and mature myeloid, lymphoid, and erythroid or megakaryocytic subsets were determined by flowcytometry using antibody panels detailed in Supplementary Table **3**. In brief, cells were isolated from engrafted mice, blocked with human/murine Fc block, and stained with indicated combinations of antibodies. Data were acquired with FACSDiva on a LSR Fortessa (BD Biosciences) equipped with 5 lasers and analyzed with FlowJo V10 software.

*Histologic Analysis:* Tissues were fixed in Bouin’s Fixative solution (RICCA Chemical Company, TX, USA) and embedded in paraffin. Femurs were decalcified with Formic Acid Bone Decalcifier (Decal Chemical, NY, USA). Paraffin blocks were sectioned at 4 μm and stained with hematoxylin and eosin (H&E) or antigen-specific antibodies routinely used in the Yale Clinical Pathology and Yale Pathology Tissue Services (**Supplementary Table 4**). Images were acquired using Nikon Eclipse 80i microscope.

All animal experimentations were performed in compliance with Yale Institutional Animal Care and Use Committee protocols.

### MSC Culture and Verification of Immunophenotype and Human Cytokine Expression

Human or mouse BM was cultured in Dulbecco's Modified Eagle Medium (DMEM, ThermoFisher, Wilmington, DE) supplemented with 20%FBS, Pen/Strep, and L-Glutamine. Non-adherent cells were removed after 24 hours of culture and MSC grown to confluency. Cells were expanded for a maximum of 5 passages and their immunophenotype confirmed by flow cytometry using antibody panels detailed in **Supplementary Table 3** and analyzed on a LSR Fortessa. Total RNA was isolated using the RNeasy Mini Kit (Qiagen, Valencia, CA) and reverse-transcribed with the ISCRIPT™ cDNA Synthesis kit (Bio-Rad). Primers for verification of cytokine expression are provided in **Supplementary Table 5**. Reaction products were analyzed on 2%agarose gels. Q- RT-PCR was run in triplicates per standard protocol using the same primers.

### Conventional Karyotyping and FISH

Conventional chromosome analysis was performed on flow-sorted human CD45+ cells from mouse BMs according to the Yale Clinical Genetics Laboratory standardized protocols. Routine FISH tests were performed using a MDS/AML panel of commercial probes for the 5q (*EGR1* gene at 5q31), 7q (*D7S486* at 7q31), 8q (*MYC* at 8q24) and 20q (*D20S108* at 20q12) loci (Abbott Molecular, Des Plaines, IL).

### Targeted Exome Capture, Sequencing, and Analysis

DNA was digested using the QIAamp DNeasy blood and tissue DNA extraction kits (Qiagen), according the manufacturer’s recommendations. Purity and concentration of the extracted DNA was measured using NanoDrop 1000 spectrophotometer (Thermo Scientific) and Quant-iT PicoGreen dsDNA Assay Kit (Life Technologies, Carlsbad, CA) for all samples.

A library of coding exons and intron-exon boundaries of 142 genes (see Supplemental Table **6**) known to carry mutations in myeloid malignancies and cancers was prepared using the HaloPlex target enrichment kits and HaloPlex HS Target Enrichment System (Agilent Technologies, Santa Clara, CA) according to manufacturer’s instructions. In brief, approximately 200ng of DNA was fragmented using restriction enzymes proprietary to the kit. For mixed human/mouse samples isolated from MISTRG mice total DNA (human/mouse) input was calculated with the endpoint of 200ng human DNA input based on engraftment percentage of huCD45+ muCD45^-^ cells determined by flow-cytometry. Probes with sequence indexes were hybridized to the targeted DNA fragments. Each probe is designed to hybridize to both ends of a targeted DNA restriction fragment resulting in their circularization. The biotinylated probe-DNA fragment hybrids were retrieved with magnetic streptavidin beads. Small fragments of <150bp and unligated probes were removed from the mix by AMPure purification (Agencourt Bioscience, Beverly, MA). Circular molecules were ligated and enriched DNA fragments were amplified with universal primers. Quality of the libraries was verified using the Tape Station 4200 (Agilent) and input DNA estimated using a library quantification kit (Kapa Biosystems, Wilmington, MA, USA). For samples Y013, Y014, Y016, Y019, Y021, Y022, Y028, Y029 and their engrafted NSG and MISTRG mice, a second-generation enrichment kit was used with Agilent’s improved high sensitivity technology with addition of molecular barcodes to each probe. Sequencing was performed on Illumina HiSeq 2000 using 74 base pairs paired-end reads, HiSeq 4000 using 100 base pairs paired-end reads or MiSeq using 250 base pairs paired-end reads. Reads were filtered by Illumina CASAVA 1.8.2 software, and trimmed at the 3’ end using FASTX v0.0.13. To remove potential mouse contamination, each read pair was aligned to a concatenated genome of human (GRCh37) and mouse (mm10) reference genome by Burrows-Wheeler Aligner v0.7.5a. Only read pairs that were specifically aligned to human reference genome were extracted for the downstream analysis. Local realignment was performed around putative and known insertion/deletion (INDEL) sites using RealignerTargetCreator (Genome Analysis Toolkit: GATK v3.1.1) and applied base quality recalibration using GATK. MuTect v.1.1.4 and Strelka v.1.0.14 were applied to call somatic single nucleotide variants (SNV) and indels, respectively. WES data from 10 external normal blood samples were pooled to serve as reference normal cohort for somatic variant calling by MuTect and Strelka. In each sample, low confidence somatic calls were removed by applying the following filters: (i) variants with total coverage <50, (ii) with a ratio of mutant allele frequency (MAF) in tumor versus normal <5, (iii) variant base quality < 20. Variants that were considered likely to be germline because they were listed in any of the following datasets dbSNP, ESP6500, 1000Genome or Exac01, or had MAF < 0.02 in the tumor samples were excluded from further analysis. Recurrent (N>5 cases) annotated variants in COSMIC v64 and Clinvar (http://www.ncbi.nlm.nih.gov/clinvar/) were white-listed. At last, only non-synonymous variants were kept. To extract the allele frequency of the variants, all nonsynonymous somatic mutations from all the samples associated with certain patients were aggregated. The allele frequency of each variant was assessed using Samtools mpileup.

### Cloning and Isogenic Cell Line Construction

To generate isogenic wildtype and mutant expressing cell lines the lentiviral plasmids pSLIK-IDH2-FLAG and pSLIK-IDH2-R172K-FLAG (a gift from Christian Metallo, Addgene plasmid ## 66806, 6680, ^48^) were used to transduce human erythroid leukemia cell (HEL) cell lines. To generate IDH2 R140Q mutant plasmid IDH2-WT-FLAG was amplified and cloned into pGEM®-T Easy Vector Systems (A1360, Promega, Madison, WI). IDH2 R140Q site-directed mutagenesis was preformed using the QuikChange II Site-Directed Mutagenesis Kit (Agilent Technologies) with primers provided in **Supplementary Table 5** and cloned into the pEN_TTmcs entry vector for recombination into the pSLIK-hygro lentiviral vector (kind gifts from Iain Fraser, Addgene plasmid # 25755 and # 25737, respectively, ^49^). Viral particles were produced by cotransfection of 293FT cells (Life Technologies) with psPAX2 (a gift from Didier Trono (Addgene plasmid # 12260)) and pCMV-VSVG (a gift from Tannishtha Reya (Addgene plasmid # 14888)) using lipofectamine transfection reagent (Life Technologies). HEL cells were grown in RPMI/10%FBS/1%P/S/G and transduced at a multiplicity of infection (MOI) of 1, Hygromycin selected to generate HEL/ IDH2 WT, R172K and R140Q expressing cell lines. Doxycycline-inducible expression (1mg/ml, Sigma-Aldrich, St Louis, MO) was verified by sanger sequencing and western blotting with anti-Flag antibody (Clone M2, Sigma-Aldrich).

### Enasidenib Treatment

Enasidenib was purchased from Shanghai ZaiQi Bio-Tech, China and LeadGen Labs, USA and quality verified by the Yale Center for Molecular Discovery. Cell proliferation and suppression of 2-HG production was verified *in vitro* by treating dox-induced HEL-IDH2 WT/R172K/R140Q cells and primary human leukemia cells. Primary IDH2 WT and R140Q leukemia MNC were cultured in StemSpan (Stem Cell Technologies, Vancouver, Canada) supplemented with 1%P/S and recombinant human cytokines FLT-3 (50ng/ml), SCF (50ng/ml),THPO (100ng/ml), IL-3 (10ng/ml), IL-6 (25ng/ml). All cytokines were purchased from NeoBioSci, MA, USA. Enasidenib dosing was optimized and added to cell cultures (20nM) every other day. For *in vivo* treatment enasidenib was dissolved in 0.5%methylcellulose and 0.2%Tween80 in PBS at a concentration of 4mg/mL and administered via oral gavage twice daily at 40mg/kg. 2-HG was measured in cell culture supernatants and in plasma of IDH2^WT^ and IDH2^R140Q^ leukemia engrafted MISTRG before and after treatment with enasidenib or vehicle. 2-HG was measured in triplicates with the D-2-Hydroxyglutarate (D2HG) Assay kit (Biovision, CA,USA) according to manufacturer’s protocol.

### Statistical Analysis

Data was analyzed using Prism 7 (GraphPad Software, La Jolla, CA) with the use of Fisher's exact test, unpaired t-test, unpaired Mann-Whitney U test, one-way ANOVA Kruskal-Wallis test with Dunn’s Multiple Comparison Test, or two-way ANOVA with Sidak’s Multiple Comparison Test as indicated. P value was considered significant at values less than 0.05. Linear regression analyses were performed with R. Pearson correlation p-values were determined with the cor.test () function implemented in R.

## Acknowledgements

**Acknowledgements** We thank all our patients. We thank all clinicians and clinical staff for their help with patient recruitment. We thank Autumn DiAdamo for FISH and cytogenetics services. We thank C. Weibel and G. Wylezinski for technical assistance in the animal studies. We thank A. Brooks for research histology services. We thank Drs. Diane Krause for critical reading of the manuscript. We thank G. Yancopoulos, D. Valenzuela, A. Murphy and W. Auerbach at Regeneron Pharmaceuticals who generated, in collaboration with our groups, the individual knock-in alleles combined in MISTRG. This study was supported by an Institutional Research Grant number 93-032-16 from the American Cancer Society (to S.H.), the U.S. Department of Defense Peer Reviewed Cancer Research Program Career Development Award (CA120128, W81XWH-12-PRCRP-CDA; to S.H.), the State of Connecticut Department of Public Health RFP 2014-0135 (to S.H.), the Edward P. Evans and the Frederick Deluca Foundations (to S.H.). E.M.B. is supported by the National Institutes of Health (NHLBI, R01HL123851). Y.K. is supported by the National Institutes of Health (NHGRI, R01HG008383). M.G.M. is supported by the Swiss National Science Foundation (310030_146528/1), the University of Zurich Clinical Research Priority Program Human Hemato-Lymphatic Diseases. R.A.F. and M.G.M. are supported by the Bill and Melinda Gates Foundation, and the National Institutes of Health, National Cancer Institute (grant CA156689). R.A.F. is supported by the Howard Hughes Medical Institute. This study was supported by the Animal Modeling Core of the Yale Cooperative Center of Excellence in Hematology (NIDDK U54DK106857) and the Yale Hematology Tissue Bank at Yale University School of Medicine and the Yale Comprehensive Cancer Center.

## Author Contributions

Y.S., A.R., A.T., R.G., X.W., K.B., A.B., Y.T., E.B., and S.H. designed research, performed experiments, and analyzed data. T.J., T.T., J.Z., and Y.K. analyzed exome data and provided statistical analysis. N.P., N.N., E.W., and R.T. provided patient samples, data, and clinical input. Y.S., M.X., and S.H. analyzed histology and flow-cytometry data. Y.S., T.J., M.G.M, R.A.F., and S.H. wrote the manuscript. R.A.F. and S.H. directed the study.

**Author Information** The authors declare no competing financial interests. Correspondence and requests for materials should be addressed to S.H. or R.A.F.

**Abbreviations**
PB: peripheral blood
BM: bone marrow
IHC: immunohistochemistry
H&E: hematoxylin&eosin
S.E.M.: Standard Error of the Mean
MDS-MLD: Myelodysplastic syndrome with multilineage dysplasia
MDS-RS-SLD: Myelodysplastic syndrome with ring sideroblasts with single lineage dysplasia
MDS-EB: Myelodysplastic syndrome with excess blasts
MSCs: Mesenchymal stromal cells
HSC: Hematopoietic stem cells
M-CSF: macrophage colony stimulating factor
GM-CSF: granulocyte-macrophage colony stimulating factor
THPO: Thrombopoietin

